# Role of a TonB-dependent receptor and an oxygenase in iron-dependent copper resistance in *Caulobacter crescentus*

**DOI:** 10.1101/2024.11.25.625226

**Authors:** Pauline Cherry, Hala Kasmo, Mauro Godelaine, Françoise Tilquin, Marc Dieu, Patsy Renard, Jean-Yves Matroule

## Abstract

Copper (Cu) is potentially threatening for living organisms owing to its toxicity at high concentrations, requiring the onset of diverse detoxification strategies to maintain fitness. We previously showed that the environmental conditions modulate the response of the oligotrophic alphaproteobacterium *Caulobacter crescentus* to Cu excess. In the present study, we investigated the role of the Fe-importing TonB-dependent receptor (TBDR) CciT and its partner, CciO, a 2-oxoglutarate/Fe^2+^-dependent oxygenase, in Cu resistance. CciT is specifically involved in Cu resistance in both rich and poor media. Using inductively coupled plasma optical emission spectrometry, we found that under Cu stress, the cellular Cu content is reduced by overexpression of *cciT*, while Fe content increases. Mutations of the three known Fe-importing TBDRs reveal that CciT is the primary Fe importer in these conditions and the only TBDR required for Cu resistance. In addition, the extracellular Fe concentration is positively correlated with the cellular Fe content and negatively correlated with the cellular Cu content, resulting in the protection of the cells against Cu excess. The operon organization of *cciT* and *cciO* is highly conserved across bacteria, indicating a functional link between the two proteins. Deletion of *cciT*, *cciO*, or both genes leads to similar Cu sensitivity. Catalytic mutations in CciT and CciO also result in Cu sensitivity. While CciO is not required for Cu and Fe transport, its precise function remains unknown. Overall, this study provides new insights into the role of Fe uptake in Cu resistance, emphasizing the critical influence of environmental conditions on bacterial physiology.

**Importance:** Copper is an essential metal for many living organisms, as it helps to drive crucial chemical reactions. However, when present in excess, copper turns toxic due to its high reactivity with biological molecules. Bacteria may encounter excess copper in various environments, such as polluted soils, agricultural copper treatments, and within the vacuoles of infected macrophages. In this study, we investigated the copper response in the environmental bacterium *Caulobacter crescentus*. Our findings reveal that environmental iron levels play a critical role in copper resistance, as increased iron prevents cellular copper accumulation and toxicity. We identified two essential proteins, CciT and CciO, that are involved in iron transport, providing protection against copper excess.

## Introduction

Copper (Cu) is a key metal cofactor for aerobic unicellular and multicellular organisms, where it contributes to the activity of specific redox enzymes including the superoxide dismutase and the cytochrome *c* oxidase [1]. However, bacteria can encounter high levels of Cu in various environments, such as polluted soils and the vacuoles of infected macrophages causing cellular toxicity [2]. Cu toxicity results from its ability to trigger ROS formation via a Fenton-like reaction or by destabilizing the GSH/GSSG balance [3], [4]. According to the Irving-Williams series, Cu^2+^ can displace several transition metal divalent cations from metalloproteins causing mismetalation and metalloprotein dysfunction [5]. The best-known mismetalation event is the displacement of iron (Fe) atoms from Fe-S clusters [6], [7], impairing branched amino acid and heme biosynthesis for instance [6], [8]. The fate of the Fe ions released from the Fe-S clusters and their potential involvement in ROS generation is not well understood. However, these Fe ions do not seem to be recycled intracellularly, likely causing a Fe starvation response [6], [9].

In Gram-negative bacteria, the outer membrane TonB-dependent receptors (TBDRs) play a key role in extracellular Fe³⁺ uptake throughout their plugged 22 β-sheet barrel [10]. TBDRs bind extracellular Fe³⁺-siderophore complexes for subsequent entry into the periplasm [10]. The import of these complexes is energized by the TonB-ExbBD complex binding the TBDR TonB box, triggering the partial unfolding of the plug [11], [12]. It has been suggested that the number of TBDR-encoding genes in bacterial genomes is linked to the complexity of the bacterial environment [13]. The genome of the oligotrophic alphaproteobacterium *Caulobacter crescentus* encodes 62 *tbdr* genes, 4 of which have been associated with Fe uptake [14].

We have previously shown that the environment modulates Cu response in *C. crescentus*, with a stronger impact on the transcriptome when bacteria are grown under nutrient-limiting conditions [15].

In the present study, we aimed to characterize a TBDR involved in Fe import, which is associated with a 2-oxoglutarate/Fe^2+^-dependent oxygenase in the Cu resistance of *C. crescentus*. We provide evidence highlighting the critical role of proper Fe homeostasis, facilitated by the TBDR, for effective Cu resistance. The strong connection between Fe and Cu homeostasis underscores the importance of considering crosstalk between metal regulatory systems in bacterial metal resistance.

## Results

The genetic screening of a *C. crescentus* mini-Tn5 transposon mutant library seeking Cu-sensitive mutants identified the CCNA_00028 gene as one of the best candidates, with multiple transposon insertions. This gene will be hereafter referred to as *cciT* (*<u>C</u>. <u>c</u>rescentus* iron <u>t</u>ransporter). *cciT* encodes a TonB-dependent receptor (TBDR) and its expression is upregulated under moderate Cu stress in rich (PYE) and mineral (M2G) media [15]. To validate the key role of CciT in Cu resistance, we proceeded to an in-frame deletion of the *cciT* gene in the NA1000 strain and we monitored the fitness of the resulting Δ*cciT* mutant under moderate Cu stress. Consistent with the genetic screen, the Δ*cciT* mutant displayed a growth defect in PYE or M2G liquid culture supplemented with 150 µM and 15 µM CuSO_4_, respectively (Fig. 1A), whereas no growth defect was observed under control conditions (Fig. S1). The ectopic expression of *cciT* from the low copy plasmid pMR10 under the control of the constitutive lac promoter in the Δ*cciT* mutant (Δ*cciT*+) restored the WT phenotype (Fig. 1A). This observation indicates that the Cu sensitivity of the Δ*cciT* mutant does not result from a polar effect. We also tested the growth capacity of the WT, Δ*cciT,* and Δ*cciT*+ strains on solid medium by spotting ten-fold serial dilutions of liquid cultures. The Δ*cciT* mutant displayed smaller colonies with similar CFU counting when grown on M2G plates, while no growth defect was observed on PYE plates (Fig. 1B). Consistent with the observations in liquid culture, the Δ*cciT* mutant exhibits an increased Cu sensitivity when grown on M2G or PYE plates supplemented with CuSO_4_ (Fig. 1B). The Cu sensitivity returns to the WT level in the Δ*cciT*+ strain.

**Figure 1.**
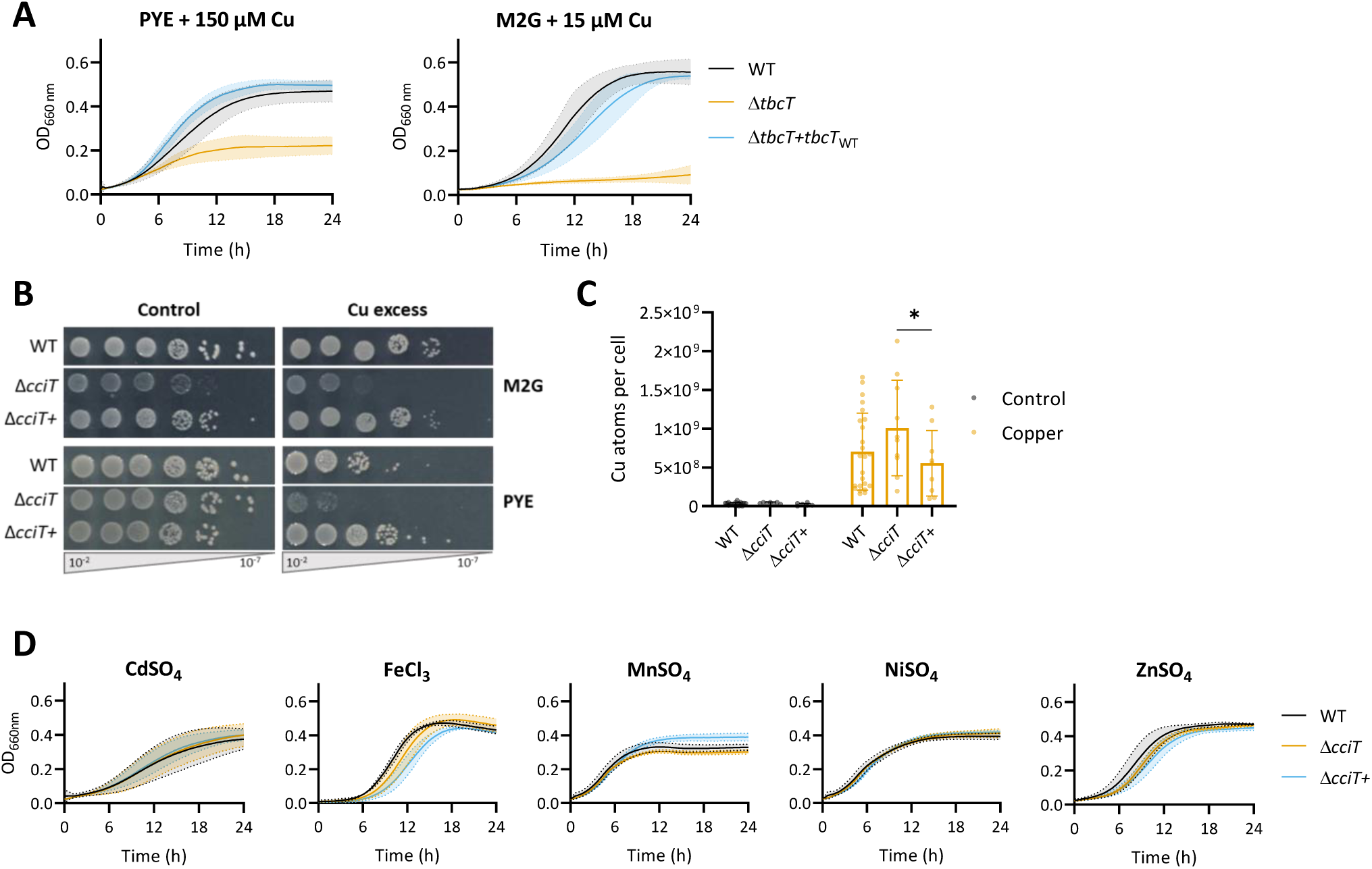
Copper resistance is relying on a TonB-dependent receptor. **A.** Growth profiles at an absorbance of 660 nm of WT, Δ*cciT*, and Δ*cciT*+ strains in PYE (top) and in M2G (bottom) media supplemented with CuSO_4_. Mean ± SD, at least three biological replicates. **B.** Viability assay on M2G and PYE plates of WT, Δ*cciT*, and Δ*cciT*+ strains, in control and CuSO_4_ excess conditions. **C**. Number of Cu atoms per cell in control condition and exposed to CuSO_4_ excess for 5 min. Mean ± SD, at least eight biological replicates. *p* values were calculated using ANOVA combined with Tukey multiple comparison test (\**p* < 0.05). **D.** Growth profiles at an absorbance of 660 nm of WT, Δ*cciT*, and Δ*cciT*+ strains in PYE exposed to 6 µM CdSO_4_, 500 µM FeCl_3_, 800 µM MnSO_4_, 200 µM NiSO_4_, 75 µM ZnSO_4_. Mean ± SD, at least three biological replicates.

Considering the role of TBDRs in metal transport, we wondered whether the Cu sensitivity of the Δ*cciT* mutant could result from a defect in Cu homeostasis. Therefore, we measured the intracellular Cu content under control and Cu stress conditions by inductively coupled plasma optical emission spectrometry (ICP-OES). The Cu content of the Δ*cciT* mutant was below the detection threshold when the bacteria were grown in M2G (Fig. 1C, left) [3], [16]. A 5 min exposure to 15 µM CuSO_4_ led to a significant increase of the cellular Cu content in all tested genetic backgrounds, but no significant difference could be observed between the WT and the Δ*cciT* strains. However, the Δ*cciT*+ strain seems to accumulate less Cu than the Δ*cciT* mutant (Fig. 1C, right). These observations indicate that the Cu sensitivity of the Δ*cciT* mutant does not result from an enhanced Cu accumulation.

To assess the potential role of CciT in the resistance to other metal stresses, we grew the Δ*cciT* mutant in liquid PYE medium supplemented with cadmium, Fe, manganese, nickel, and zinc, which are known to trigger mismetalation events and/or oxidative stress like Cu [3], [4], [5], [17]. Surprisingly, *cciT* deletion did not increase the sensitivity of *C. crescentus* to these metals (Fig. 1D), suggesting that CciT is specifically dedicated to Cu resistance.

TBDRs are mostly known for their role in the uptake of Fe^3+^-bound siderophores [10]. An *in-silico* analysis of the CciT protein sequence reveals a 30.41 %, 41.37 %, and 38.30 % similarity with *Escherichia coli* Fiu TBDR and with *Acinetobacter baumanii* and *Pseudomonas aeruginosa* PiuA TBDR, respectively. Fiu and PiuA have been described as catechol-derived siderophore importers [8–10]. In *C. crescentus*, the expression of the *cciT* gene is under the control of the ferric uptake regulator Fur and is upregulated under Fe-limiting conditions [20]. To assess the role of CciT in Fe homeostasis, we measured the cellular Fe content of the Δ*cciT* mutant by ICP-OES. The Δ*cciT* mutant harbors a similar Fe content to the WT strain, while the overexpression of *cciT* in the Δ*cciT*+ strain (Fig. S2) leads to an increased Fe accumulation relative to the Δ*cciT* mutant (Fig. 2A). These data are in line with the role of CciT in Fe import and a potential redundancy with other Fe importers. Consistent with the latter hypothesis, the Δ*cciT* mutant displays a WT-like growth profile in Fe-limiting liquid medium (Fig. 2B). Yet, the dispensability of CciT under Fe-limiting conditions is less obvious on solid Fe-depleted M2G medium, where the Δ*cciT* mutant exhibits a strong growth defect. This suggests that the access to Fe in liquid and solid media is different. Normal growth is restored in the Δ*cciT*+ strain or by Fe repletion in the medium (Fig. 2C), reinforcing the role of CciT in Fe uptake. Interestingly, Cu treatment does not impact the cellular Fe content (Fig. 2A).

**Figure 2.**
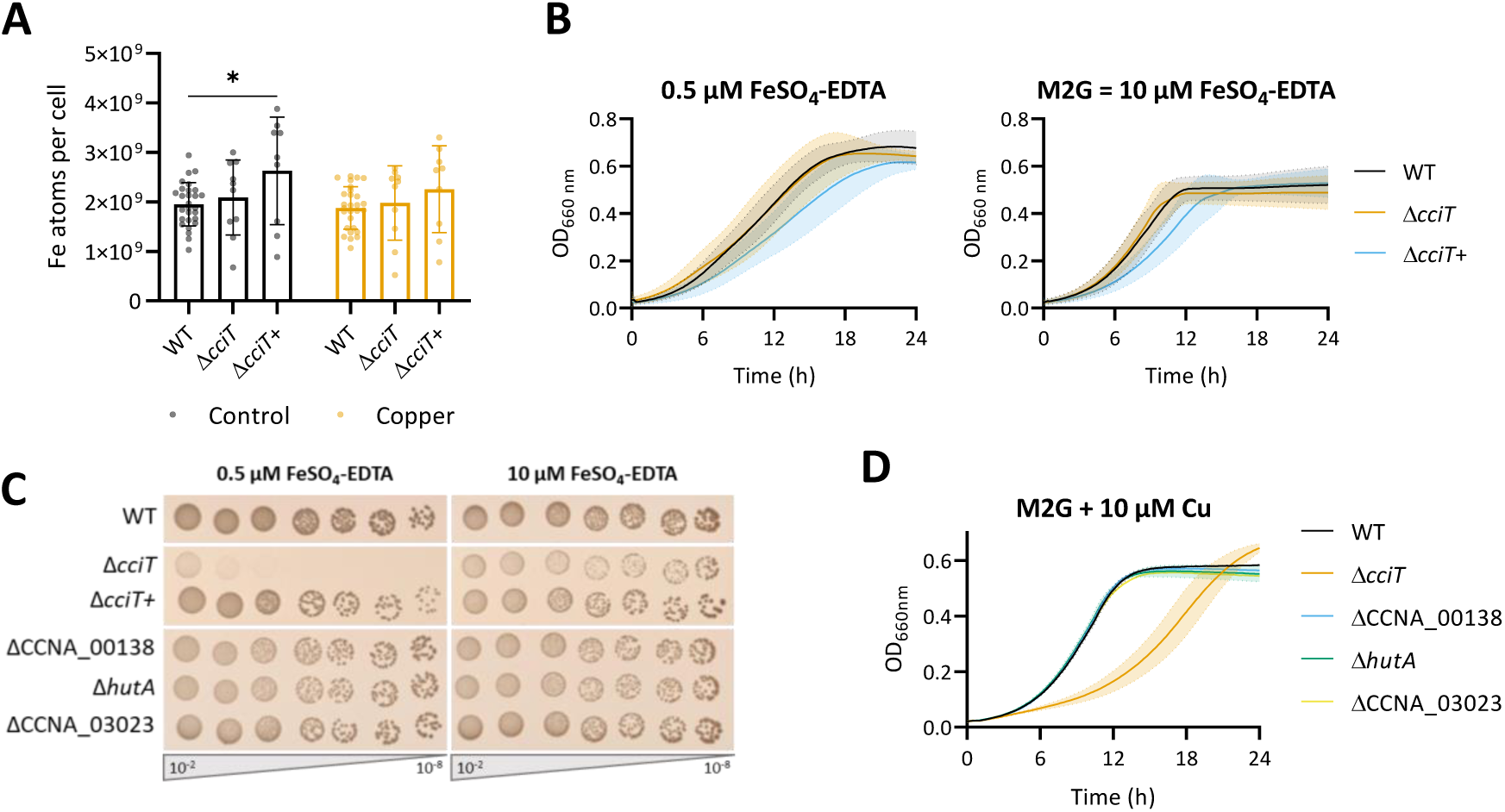
CciT is involved in Fe uptake. **A.** Number of Fe atoms per cell in control condition and exposed to CuSO_4_ excess for 5 min. Mean ± SD, at least eight biological replicates. *p* values were calculated using ANOVA combined with Tukey multiple comparison test (\**p* < 0.05). **B.** Growth profiles at an absorbance of 660 nm of WT, Δ*cciT*, and Δ*cciT*+ strains in M2G medium in Fe-limiting (left) and Fe-replete (right) conditions. Mean ± SD, at least three biological replicates. **C**. Viability assay on M2G plates of WT, Δ*cciT*, Δ*cciT*+, ΔCCNA_00138, Δ*hutA* and ΔCCNA_03023 strains, in Fe-limiting (left) and Fe-replete (right) conditions. **D.** Growth profiles at an absorbance of 660 nm of WT, Δ*cciT*, ΔCCNA_00138, Δ*hutA* and ΔCCNA_03023 strains in M2G media exposed to CuSO_4_ excess for 5 min. Mean ± SD, at least three biological replicates.

The genome of *C. crescentus* harbors 61 additional TBDR-encoding genes, among which CCNA_00138, *hutA,* and CCNA_03023 genes have been associated with Fe homeostasis, owing to their Fur-dependent expression under Fe-limiting conditions [20]. To validate their potential role in Fe homeostasis, we generated the ΔCCNA_00138, Δ*hutA,* and ΔCCNA_03023 knock-out mutants by in-frame deletion and measured their growth on solid M2G medium under limiting Fe conditions, where the Δ*cciT* mutant showed a growth defect. None of these 3 mutants was affected by Fe limitation, suggesting that CciT could be the main Fe importer in these conditions (Fig. 2C). The expression of the CCNA_00138, *hutA,* and CCNA_03023 genes is upregulated under moderate Cu stress [15], suggesting that they may play a role in Cu resistance. Yet, none of the ΔCCNA_00138, Δ*hutA,* and ΔCCNA_03023 mutants exhibited an increased Cu sensitivity (Fig. 2D). This observation reinforces the central role of CciT in Cu resistance, possibly through Fe import.

To explore the link between optimal Fe uptake and Cu resistance in *C. crescentus*, we monitored the Cu sensitivity of the WT strain at 2.5 μM, 10 µM, and 50 μM FeSO_4_-EDTA corresponding to low, standard, and high Fe concentrations, respectively. The low and high Fe conditions do not seem to impact bacterial fitness under control conditions (Fig. 3A, black curve). However, under Cu stress, the extracellular Fe concentration correlates with the extent of Cu tolerance, ranging from an absence of bacterial growth under low Fe to a loss of Cu sensitivity at high Fe concentration (Fig. 3A, orange curve). In support of these observations, the cellular Cu content measured by ICP-OES negatively correlates with the extracellular Fe concentration (Fig. 3B). One may argue that the high concentration of the broad metal chelator EDTA used in high Fe conditions leads to Cu chelation in the extracellular medium, preventing its entry and explaining the reduced cellular Cu content in this condition (Fig. 3B, left panel). To test this hypothesis, we assessed the proteome of the WT strain using LC-MS on bacteria grown in low, standard, and high Fe conditions. We monitored the abundance of the stress response CpxP protein (CCNA_03997), which accumulates when *C. crescentus* is exposed to Cu excess. The striking increase in abundance of CpxP in all tested Fe conditions (Fig. 3C) suggests that Cu entered the cells, at least long enough to trigger a Cu stress. One could then hypothesize that an unknown Cu efflux system is activated when extracellular Fe concentration is increasing. Collectively, these data support a tight link between Cu homeostasis and Fe homeostasis.

**Figure 3.**
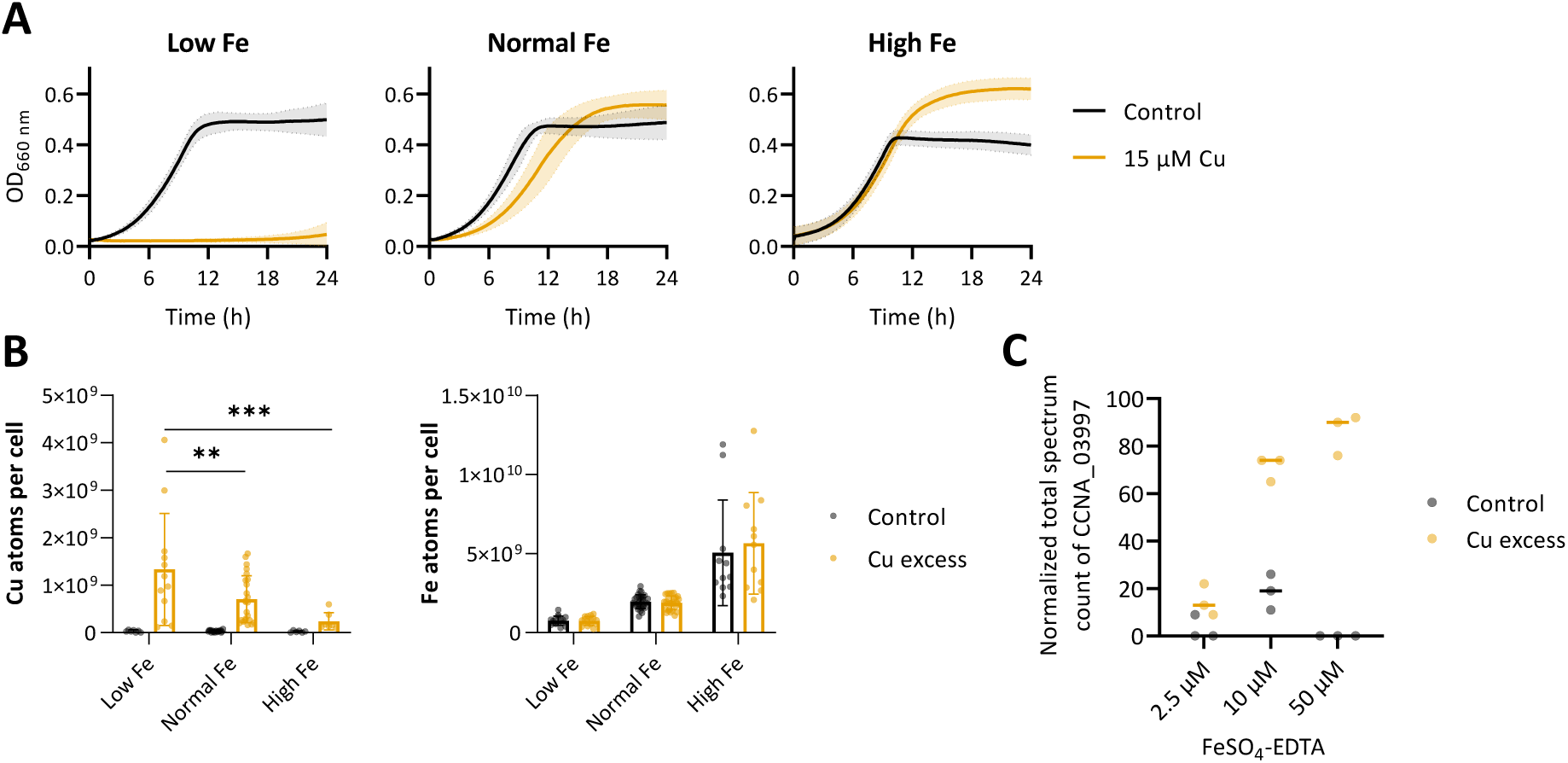
Intracellular Fe content supports Cu resistance in *C. crescentus*. **A.** Growth profiles at an absorbance of 660 nm of WT strain in modified M2G medium, with low, normal and high Fe concentrations, combined to CuSO_4_ excess. Mean ± SD, at least three biological replicates. **B.** Number of Cu (left) and Fe (right) atoms per cell in control condition and exposed to 5 minutes CuSO_4_ excess. Mean ± SD, at least nine biological replicates. *p* values were calculated using ANOVA combined with Tukey multiple comparison test (\**p* < 0.05, \*\**p* < 0.01, \*\*\**p* < 0.005). **C.** Normalized total spectrum count of CCNA_03997 protein in low, normal and high Fe conditions, in combination with a 1h exposure to CuSO_4_ excess.

The CciT-encoding gene is part of an operon with the CCNA_R0097 sRNA gene, and the CCNA_00027 gene, predicted to encode a 2-oxoglutarate/Fe^2+^-dependent oxygenase (2OGX), hereafter referred to as CciO (Fig. 4A) [21]. A similar genomic organization is observed in *E. coli*, where the *fiu* gene is in an operon with the 2OGX-encoding *ybiX* and *ybiI* coding for a Zn finger domain-containing protein [22]. In *P. aeruginosa* and *A. baumanii*, *piuA* is located next to the PiuC 2OGX-encoding gene [18]. To determine whether this cooccurrence can be extended to other bacterial species, we proceeded to synteny search using the FlaGs bioinformatic tool, which clusters the 4 neighboring genes of the up-and downstream regions of the gene of interest [23]. The database was constructed by compiling the orthologs of CciO and CciT through the KEGG orthology groups [24]. The genes encoding the protein homologs of CciT and CciO are reciprocal best neighbors as 87.7 % of CciO-encoding genes have a neighboring CciT-encoding gene, and 68.2 % of examined CciT-encoding genes have a neighboring CciO-encoding gene (Fig. 4B). The second and third best matches for CciT and CciO are a PepSY associated TM helix domain-containing protein and a Sel1 repeat family protein, respectively. Nevertheless, the two genes are in the vicinity of either *cciO* or *cciT* in less than 25 % of the genomes. When monitoring the cooccurrence of *cciT* and *cciO* genes, most genomic organizations have no insertion between CciT and CciO homologs (Fig. 4C). It also appears that CciO-homologs genes are most often found downstream the CciT homologs genes and vice-versa when considering the CciT-homologs.

**Figure 4.**
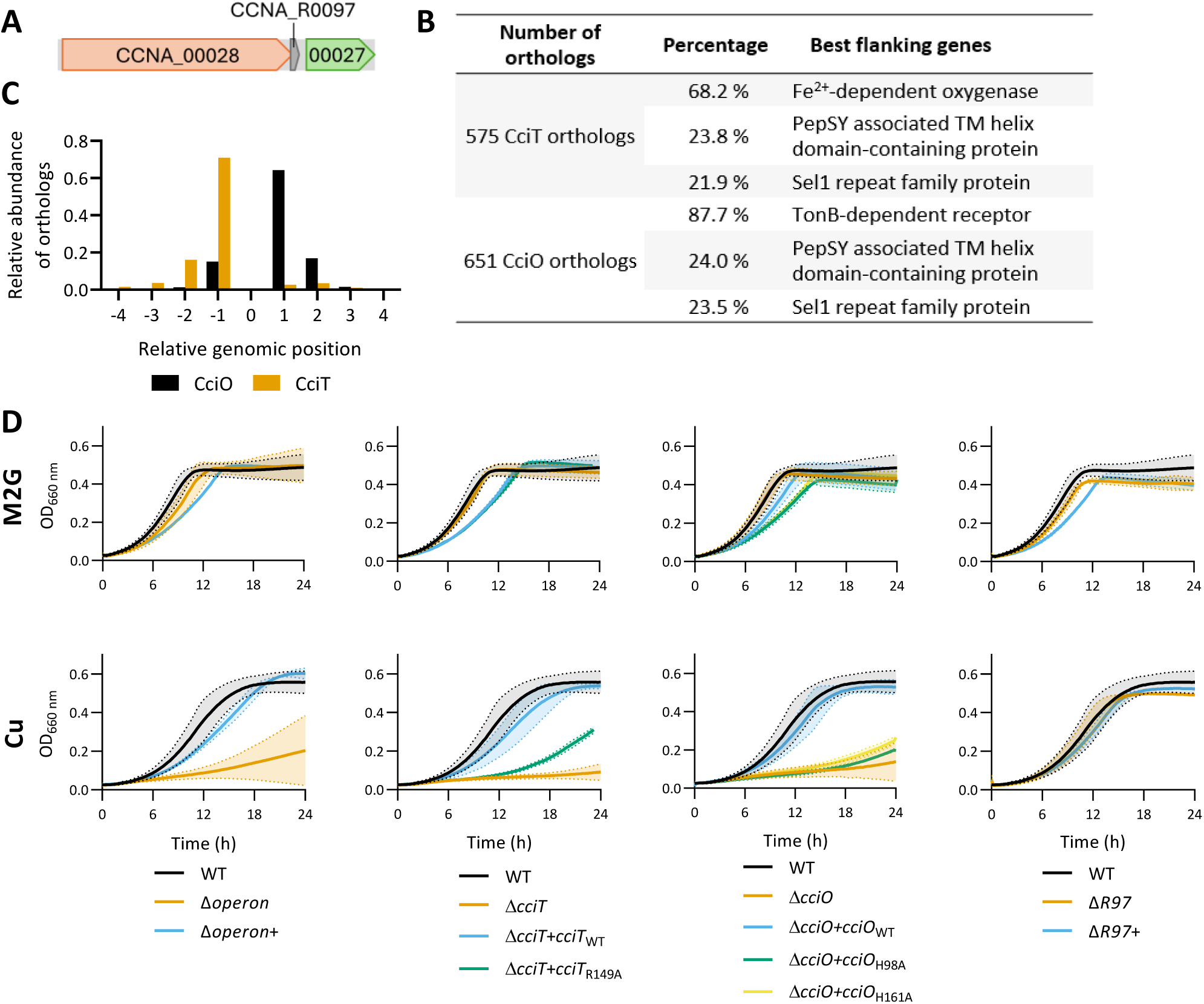
CciT and CciO operon organization is conserved. **A.** Genome organization of *cciT* (CCNA_00028), CCNA_R0097 and *cciO* (CCNA_00027) genes in *C. crescentus* NA1000. **B.** Top three clusters of protein-encoded genes identified by FlaGs in the 4 genes up-and downstream of the considered orthologs. **C.** Relative position of CciT or CciO-encoding genes, to the considered ortholog, CciO or CciT respectively. **D.** Growth profiles at an absorbance of 660 nm of WT, Δ*operon*, Δ*cciT*, Δ*cciO* and ΔR97 strains alongside respective complementation strains, in M2G medium (upper) and exposed to CuSO_4_ excess (lower). Mean ± SD, at least three biological replicates.

To determine the potential involvement of CciO and R97 in Cu resistance, we generated in-frame deletions of the *cciO* and R97 genes together with a deletion of the whole operon, and we monitored the Cu sensitivity of the resulting mutants in M2G. While the ΔR97 mutant exhibits a WT phenotype in the tested conditions, the Δ*cciO* and Δ*operon* mutants display the same Cu sensitivity as the Δ*cciT* mutant (Fig. 4D). In line with the Δ*cciT+* strain, the Cu sensitivity phenotype of the Δ*cciO* and Δ*operon* mutants is restored to the WT level by the ectopic expression of either *cciO* or the operon, respectively (Fig. 4D, Fig. S2). Based on structure prediction of CciT and CciO by Alphafold and resolved structures of their homologs (Fig. S3, S4) [19], [25], we generated point mutants of CciT and CciO to understand whether the presence and/or the function of CciT and CciO supports Cu resistance. The expression of the CciT_R149A_, CciO_H98A_, and CciO_H161A_ variants in the respective Δ*cciT* or Δ*cciO* mutants does not complement their Cu sensitivity (Fig. 4D). This observation indicates that both the presence and the activity of CciT and CciO are required to provide a complete Cu resistance, hinting towards a functional link between CciT and CciO.

We investigated the role of the CciO protein in Cu and Fe homeostasis by measuring the intracellular Cu and Fe levels in the Δ*cciO* and Δ*cciO*+ strains under both control and Cu excess conditions (Fig. 5A and B). Both strains accumulated Cu to the same extent as the WT strain under Cu excess, with *cciO* overexpression having no impact on Cu accumulation (Fig. 5A). As previously noted, the Δ*cciT*+ strain accumulates more Fe compared to Δ*cciT* under control conditions. In contrast, neither the deletion nor the overexpression of *cciO* affects the Fe content relative to the WT strain in both control and Cu excess conditions (Fig. 5B). However, the Δ*cciT*+ strain contains more Fe per bacterium than both Δ*cciO* and Δ*cciO*+ strains.

**Figure 5.**
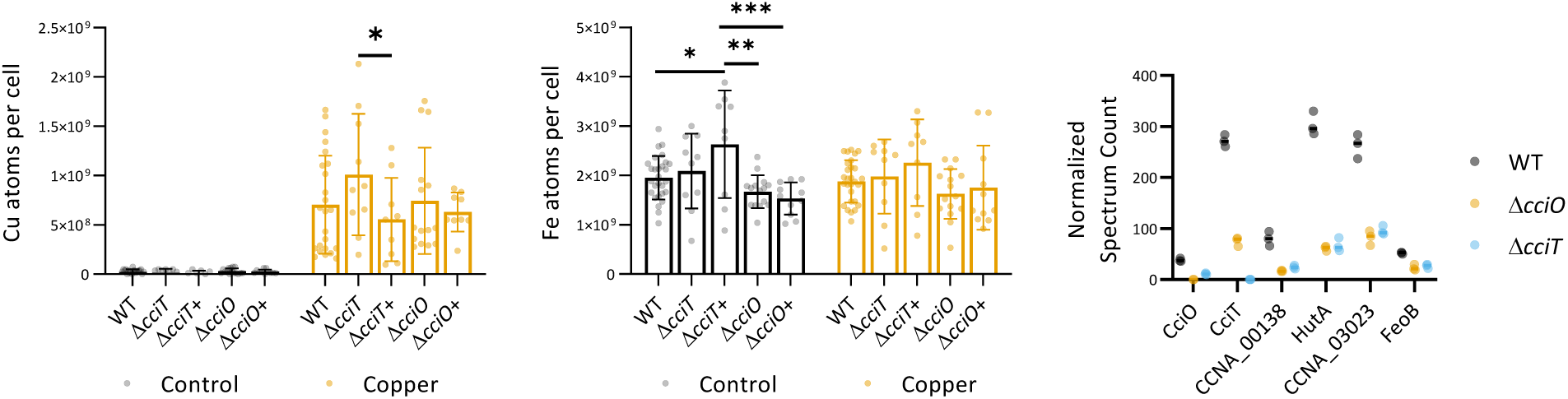
CciO is not involved in Fe uptake. **A.** Number of Cu atoms per cell in control condition and exposed to CuSO_4_ excess for 5 min. Mean ± SD, at least eight biological replicates. *p* values were calculated using ANOVA combined with Tukey multiple comparison test (\**p* < 0.05). **B.** Number of Fe atoms per cell in control condition and exposed to CuSO_4_ excess for 5 min. Mean ± SD, at least eight biological replicates. *p* values were calculated using ANOVA combined with Tukey multiple comparison test (\**p* < 0.05). **C.** Normalized spectrum count of peptides in WT, Δ*cciO* and Δ*cciT* strains grown in M2G medium, measured by LC-MS. Individual values and means represented.

To better understand the role of CciT and CciO in *C. crescentus*, we examined how the deletion of either *cciT* or *cciO* gene affects the cellular proteome, anticipating potential changes in the abundance of interacting partners as a compensatory adaptation. For this purpose, we measured the protein abundance of the Δ*cciT* and Δ*cciO* mutants using untargeted proteomics. Surprisingly, the protein levels of CciT and CciO were reduced in the Δ*cciO* and Δ*cciT* strains, respectively (Fig. 5C). This decrease cannot be attributed to polar effects from the deletion of either the *cciO* or *cciT* genes on the remaining genes within the operon, as the levels of *cciT* and *cciO* mRNA assessed by RT-qPCR remain unchanged in the Δ*cciO* and Δ*cciT* mutants (Fig. S2). We also observed a decrease in the abundance of Fe-importing TBDRs—CCNA_00138, HutA, and CCNA_03023—as well as the FeoB protein in both Δ*cciO* and Δ*cciT* strains, even though intracellular Fe levels remain unchanged in these mutants.

## Discussion

The significance of maintaining proper Fe homeostasis in Cu resistance is an emerging concept in bacteria. Our study emphasizes the role of a TonB-dependent receptor (TBDR), CciT, along with its partner CciO, a 2-oxoglutarate/Fe^2+^-dependent oxygenase (2OGX), in conferring Cu resistance in the free-living alphaproteobacterium *Caulobacter crescentus*.

In this study, we demonstrated that CciT, an Fe-importing TBDR, is essential for effective Cu resistance. While TBDRs are typically known for their role in actively importing extracellular substrates, there are exceptions—such as PopC from *Myxococcus xanthus*, which is involved in the export of the protease PopC [10], [26]. In *C. crescentus*, CciT appears to play a unique role in Cu resistance by preventing the intracellular accumulation of Cu, a function not commonly associated with TBDRs.

Previous studies proposed that CciT, along with the CCNA_00138, HutA, and CCNA_03023 TBDRs are involved in Fe homeostasis owing to their upregulation under Fe-limiting conditions [14], [20]. Consistent with these observations, we demonstrate that CciT is involved in Fe uptake, even though CciT is not essential in liquid medium under Fe-limiting conditions. One could propose that this dispensability results from redundancy with the other Fe-importing TBDRs, compensating for the absence of CciT to ensure proper Fe homeostasis. HutA has been shown to import Fe even though no siderophore biosynthesis pathway could be found in *C. crescentus* [14]. This questions the nature of the Fe^3+^-bound substrate under laboratory conditions where *C. crescentus* is the sole bacterial species in the culture. The specific requirements of CciT for proper growth on solid medium under low Fe and high Cu conditions suggest that CciT could be the main Fe importer in *C. crescentus*, at least under the tested conditions.

We also demonstrated that the environmental Fe concentration influences the intracellular Fe content of *C. crescentus*, as well as its capacity to cope with Cu excess. We observed that the environmental Fe concentration directly impacts the intracellular Fe content, which is negatively correlated with the Cu accumulation. It has been shown that Cu-exposed bacteria, such as *E. coli* and *Rubrivivax gelatinosus*, rely on Fe-uptake systems to tolerate Cu [8], [10]. The toxicity of Cu results from the mismetalation of Fe-containing proteins as well as from the degradation of Fe-S clusters [6]. Adequate Fe homeostasis and Fe content could compensate for the mismetalation and enhance the biosynthesis/repair of Fe-S clusters, leading to the increased Cu resistance in Fe-rich conditions. This could explain the role of CciT as the main Fe importer in *C. crescentus*, providing Cu resistance.

Interestingly, *cciT* is part of an operon with the *cciO* gene, which encodes a 2OGX. The similar phenotypes observed in the single mutants of *cciT* and *cciO*, along with the high conservation of this operon across various bacterial species, suggest that these two proteins work together to provide Cu resistance.

The role of CciO in Cu resistance remains to be elucidated as the protein does not seem to play a role in Fe or Cu transport. In *P. aeruginosa*, it has been proposed that the CciO homolog PiuC metabolizes the siderophore monosulfactam imported by the TBDR PiuA [28]. In *E. coli*, one SNP in the CciO homolog *ybiX*, triggering translation termination, promotes resistance to a bacteriostatic catechol-siderophore conjugate, supposedly imported by Fiu [29]. Given the conservation of this system in both *E. coli* and *P. aeruginosa*, it is plausible that CciO metabolizes the substrate imported by CciT (Fig. 6A). Another possibility is that the Fe ions imported by CciT primarily support CciO function by supplying the essential Fe^2+^ cofactor required for its activity (Fig. 6B). In turn, the unknown activity of CciO could play a crucial role in conferring Cu resistance to *C. crescentus*.

**Figure 6.**
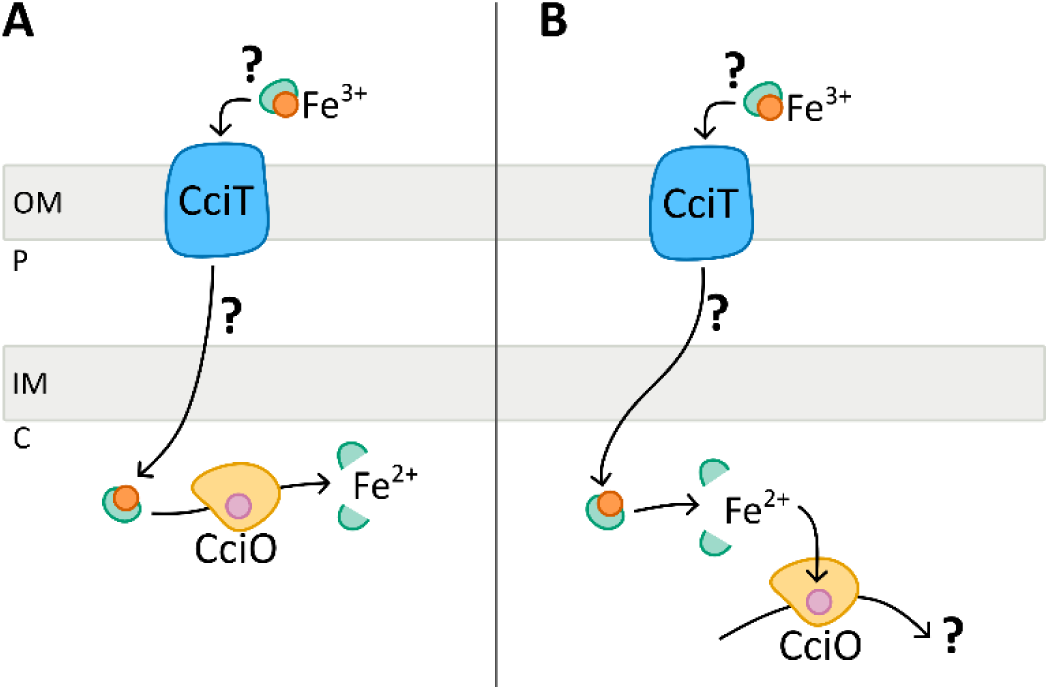
Hypothetical models for Cu resistance provided by CciT-CciO system in *C. crescentus*. **A**. CciT transports extracellular Fe^3+^ through the outer membrane via unknown carrier. The Fe^3+^ complex would be further imported into the cytoplasm, where CciO could catalyze the release of the reduced Fe^2+^ in the cytoplasm. The transport and the bioavailability of Fe would enhance Cu resistance of *C. crescentus*. **B.** The second working model considers Fe^3+^ uptake by CciT to properly metalate CciO. In turn, the unknown activity of CciO would provide Cu resistance to *C. crescentus*.

Our study revealed a strong functional link between the activity of the TBDR CciT and the 2OGX CciO, both of which play a key role in maintaining Fe homeostasis in *C. crescentus* to support effective Cu resistance. Beyond the well-known active strategies used against Cu toxicity, our findings emphasize the crucial role of environmental conditions in how bacteria cope with stress, such as Cu excess. This underscores the importance of considering the complexity of bacterial environments when investigating their physiological processes.

## Material and Methods

### Bacterial strains, plasmids and growth conditions

*Caulobacter crescentus* NA1000 was routinely grown at 30 °C, under moderate shaking, in either PYE rich medium (0.2 % bacto peptone, 0.1 % yeast extract, 1 mM MgSO_4_, 0.5 mM CaCl_2_) or mineral M2G medium (5.4 mM Na_2_PO_4_, 4 mM KH_2_PO_4_, 9.35 mM NH_4_Cl, 0.2 % w/v glucose, 10 μM FeSO_4_-EDTA, 0.5 mM MgSO_4_, 0.5 mM CaCl_2_) [263]. The media was supplemented with 5 μg/ml kanamycin, 2.5 μg/ml oxytetracycline, 15 μg/ml nalidixic acid, and/or CuSO_4_.5H_2_O when required. Exponentially growth cultures were used for all experiments. Strains and plasmids used in this study are listed in Table S1. The strategies and the primers for their construction are available upon request.

### Growth curves measurements

Bacterial cultures in exponential growth phase (OD_660 nm_ of 0.4 – 0.6) were diluted in PYE or M2G medium to a final OD_660 nm_ of 0.05 and inoculated in 96-well plates with appropriate concentration of CuSO_4_, FeSO_4_, ZnSO_4_, CdSO_4_, NiSO_4_, or MnSO_4_ when required. OD_660 nm_ was recorded every 10 min for 24 h at 30 °C under continuous shaking in an Epoch 2 absorbance reader (Biotek Instruments Inc.). When required, the concentration in FeSO_4_-EDTA of M2G medium was adapted.

### Viability assay

Overnight cultures in stationary phase were diluted in 1:10 serial dilutions up to 10^-8^. Drops of 5 μl of each dilution were spotted on PYE or M2G plates, containing CuSO_4_ and/or FeSO_4_ appropriate concentrations when required. Plates were incubated at 30 °C for 48 h and pictures were taken with Amersham Imager 600 (GE Healthcare Life Sciences).

### Determination of metal content

*C. crescentus* cells were fixed for 20 min in 2% paraformaldehyde at 4°C and then washed three times with an ice-cold wash buffer (10 mM Tris–HCl [pH 6.8] and 100 μM EDTA). Cells were resuspended in MilliQ water, and lyzed under 2.4 kbar by using a cell disrupter (Cell Disruption System, one shot model, Constant). Cell debris were removed by centrifugation at 10,000 x g for 15 min, and cell lysates were diluted in 1% HNO_3_. Samples were finally analyzed by ICP-OES with an Optima 8000 ICP-OES from PerkinElmer. As described in [2–3], cellular metal concentrations were calculated using the following formula:

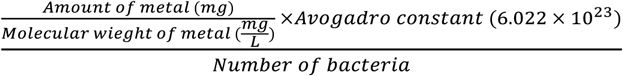

### LC-MS

Cells from 15 ml cultures of *C. crescentus* were isolated by centrifugation at 8000 rpm for 10 min at 4°C and washed three times with ice-cold wash buffer (10 mM Tris-HCl pH 6.8, 100 μM EDTA). Normalized pellets were resuspended in 2 ml of complete EDTA-free protease inhibitor cocktail (Roche, Mannheim, Germany). Cells were lyzed under 2.4 kbar by using a cell disrupter (Cell Disruption System, one shot model, Constant). For proper dissolution of membrane proteins, lysates are incubated with 0.01 % SDS for 30 min. at room temperature. Cell debris were removed by centrifugation at 19,000 x g for 15 min. The samples were treated using the optimized Filter-aided sample preparation (FASP) protocol. Briefly, the samples were loaded onto Millipore Microcon 30 MRCFOR030 Ultracel PL-30 filters that have been rinsed and washed beforehand with, 1 % Formic Acid (FA) and 8 M urea buffer (8 M urea in 0.1 M Tris buffer at pH 8.5), respectively. The proteins on the filter were then exposed to a reducing agent (dithiothreitol (DTT)) and then alkylated with iodoacetamide (IAA). The proteins were then finally digested overnight with trypsin. The final step of the digestion is to transfer proteins in 20 μl of 2% ACN (acetonitrile) and 0.1% FA, in an injection vial for inverted phase chromatography. The digest was analyzed using nano-LC-ESI-MS/MS tims TOF Pro (Bruker, Billerica, MA, USA) coupled with an UHPLC nanoElute (Bruker).

Peptides were separated by nanoUHPLC (nanoElute, Bruker) on a 75 μm ID, 25 cm C18 column with integrated CaptiveSpray insert (Aurora, ionopticks, Melbourne) at a flow rate of 400 nl/min, at 50°C. LC mobile phases A was water with 0.1% formic acid (v/v) and B ACN with formic acid 0.1% (v/v). Samples were loaded directly on the analytical column at a constant pressure of 800 bar. The digest (1 µl) was injected, and the organic content of the mobile phase was increased linearly from 2% B to 15 % in 22 min, from 15 % B to 35% in 38 min, from 35% B to 85% in 3 min Data acquisition on the tims TOF Pro was performed using Hystar 5.1 and timsControl 2.0. tims TOF Pro data were acquired using 100 ms TIMS accumulation time, mobility (1/K0) range from 0.6 to 1.6 Vs/cm². Mass-spectrometric analysis was carried out using the parallel accumulation serial fragmentation (PASEF) acquisition method [30]. One MS spectra followed by ten PASEF MSMS spectra per total cycle of 1.1 s.

All MS/MS samples were analyzed using Mascot (Matrix Science, London, UK; version 2.8.1). Mascot was set up to search the *C. crescentus* NA1000_190306 database from UniRef 100 and Contaminants_20190304 database assuming the digestion enzyme trypsin. Mascot was searched with a fragment ion mass tolerance of 0.050 Da and a parent ion tolerance of 15 PPM. Carbamidomethyl of cysteine was specified in Mascot as a fixed modification. Oxidation of methionine and acetyl of the n-terminus were specified in Mascot as variable modifications.

Scaffold (version Scaffold_5.1.1, Proteome Software Inc., Portland, OR) was used to validate MS/MS based peptide and protein identifications. Peptide identifications were accepted if they could be established at greater than 97.0% probability to achieve an FDR less than 1.0% by the Percolator posterior error probability calculation [31]. Protein identifications were accepted if they could be established at greater than 50.0% probability to achieve an FDR less than 1.0% and contained at least 2 identified peptides. Protein probabilities were assigned by the Protein Prophet algorithm [32]. Proteins that contained similar peptides and could not be differentiated based on MS/MS analysis alone were grouped to satisfy the principles of parsimony. Proteins sharing significant peptide evidence were grouped into clusters.

### RT-qPCR

Bacteria were grown in M2G up to OD_660nm_ = 0.4 before incubation with or without 15 µM CuSO_4_ for 10 min at 30 °C under agitation. Bacteria were recovered by centrifugation and pellets were flash frozen until resuspension in 40 μl of a 20 mg/ml proteinase K solution (Avantor®, Radnor, PA, USA) with 1 μl of undiluted Ready-Lyse Lysozyme solution (Lucigen®, Middlesex, UK), and lysis was allowed to proceed for 10 min in a shaking incubator at 37 °C and 600 rpm. Total RNA was retrieved from the cell suspensions using TriPure isolation reagent and procedure as described by the manufacturer (Roche, Mannheim, Germany).

RNA (2 μg) isolated from *C. crescentus* was incubated with DNAse I (Thermo Scientific, Merelbeke, Belgium) for 30 min at 37 °C. DNAse I was then inactivated with 50 mM EDTA for 10 min at 65 °C. Subsequently, RNA was subjected to reverse transcription using MultiScribe Reverse Transcriptase (Applied Biosystems®, Foster City, CA, USA) with random primers (as described by the manufacturer). A total of 300 ng of cDNA was mixed with Takyon No Rox SYBR MasterMix dTTP Blue (Eurogentec, Seraing, Belgium) and the appropriate primer sets (Supplementary Table S4) and used for qPCR in a LightCycler96 (Roche, Basel, Switzerland). Forty-five PCR cycles were performed (95 °C for 10 s, 60 °C for 10 s and 72 °C for 10 s). Primer specificity was checked by melting curves analysis. Relative gene expression levels between different samples were calculated with the 2^−ΔΔCt^ method, using the *mreB* gene as a reference. Three technical replicates were analyzed for each sample.

## Acknowledgements

We thank Valérie Charles and Carmela Aprile (CMI laboratory, NISM, UNamur) for the ICP measurements. We acknowledge Rob Van Houdt (Microbiology Unit, SCK CEN) and the URBM members for fruitful discussions.

## Author contributions

P. C. and J.-Y. M. conceptualization; P. C. methodology; P. C. validation; P. C., H. K., G. M., F. T., M. D., P. R., and J.-Y. M. formal analysis; P. C., H. K., G. M., F. T., and M. D., investigation; P. C. writing–original draft; P. C. visualization; J.-Y. M. writing–review & editing; J.-Y. M. supervision; J-.Y. M. funding acquisition.

---

**Table S1.** Bacterial strains and plasmids used in this study.

| Strains | Genotype and/or phenotype | Reference or source |
| --- | --- | --- |
| *Caulobacter crescentus* |  |  |
| WT | Wild-type NA1000, synchronizable variant of CB15 | [33] |
| $\Delta cciO$ | Knock-out strain for *cciO* gene (CCNA_00027) | This study |
| $\Delta cciO+cciO_{WT}$ | Knock-out strain for *cciO* gene carrying a copy of *cciO* on the pMR10 under the control of the lac promoter; Kan<sup>R</sup> | This study |
| $\Delta cciO+cciO_{H98A}$ | Knock-out strain for *cciO* gene carrying a copy of *cciO* with the point mutation H98A on the pMR10 under the control of the lac promoter; Kan<sup>R</sup> | This study |
| $\Delta cciO+cciO_{H161A}$ | Knock-out strain for *cciO* gene carrying a copy of *cciO* with the point mutation H161A on the pMR10 under the control of the lac promoter; Kan<sup>R</sup> | This study |
| $\Delta R97$ | Knock-out strain for the ncRNA (CCNA_R0097) | This study |
| $\Delta R97+R97$ | Knock-out strain for CCNA_R0097 ORF carrying a copy of R97 on the pMR10 under the control of the lac promoter; Kan<sup>R</sup> | This study |
| $\Delta cciT$ | Knock-out strain for *cciT* gene (CCNA_00028) | This study |
| $\Delta cciT+cciT_{wt}$ | Knock-out strain for *cciT* gene carrying a copy of *cciT* on the pMR20 under the control of the lac promoter; Tet<sup>R</sup> | This study |
| $\Delta cciT+cciT_{R149A}$ | Knock-out strain for *cciT* gene carrying a copy of *cciT* with the point mutation R149A on the pMR20 under the control of the lac promoter; Tet<sup>R</sup> | This study |
| $\Delta operon$ | Knock-out strain for *cciO* and *cciT* genes, and ncRNA R97 (CCNA_00027-R97-28) | This study |
| $\Delta operon+operon$ | Knock-out strain for *cciO* and *cciT* genes, and ncRNA R97, carrying a copy of the whole operon under the control of the lac promoter; Kan<sup>R</sup> | This study |
| $\Delta CCNA\_00138$ | Knock-out strain for CCNA_00138 gene | This study |
| $\Delta hutA$ | Knock-out strain for *hutA* gene (CCNA_02277) | This study |
| $\Delta CCNA\_03023$ | Knock-out strain for CCNA_03023 gene | This study |
| Plasmids |  |  |
| pNPTs138 | mobRP4<sup>+</sup> ori-R6K *sacB*; integrative vector in *C. crescentus* for in-frame deletions; Kan<sup>R</sup> |  |
| pMR10 | Low copy and replicative vector in *C. crescentus*; Kan<sup>R</sup> |  |
| pMR20 | Derivative of pMR10; Tet<sup>R</sup> |  |

**Figure S1.**
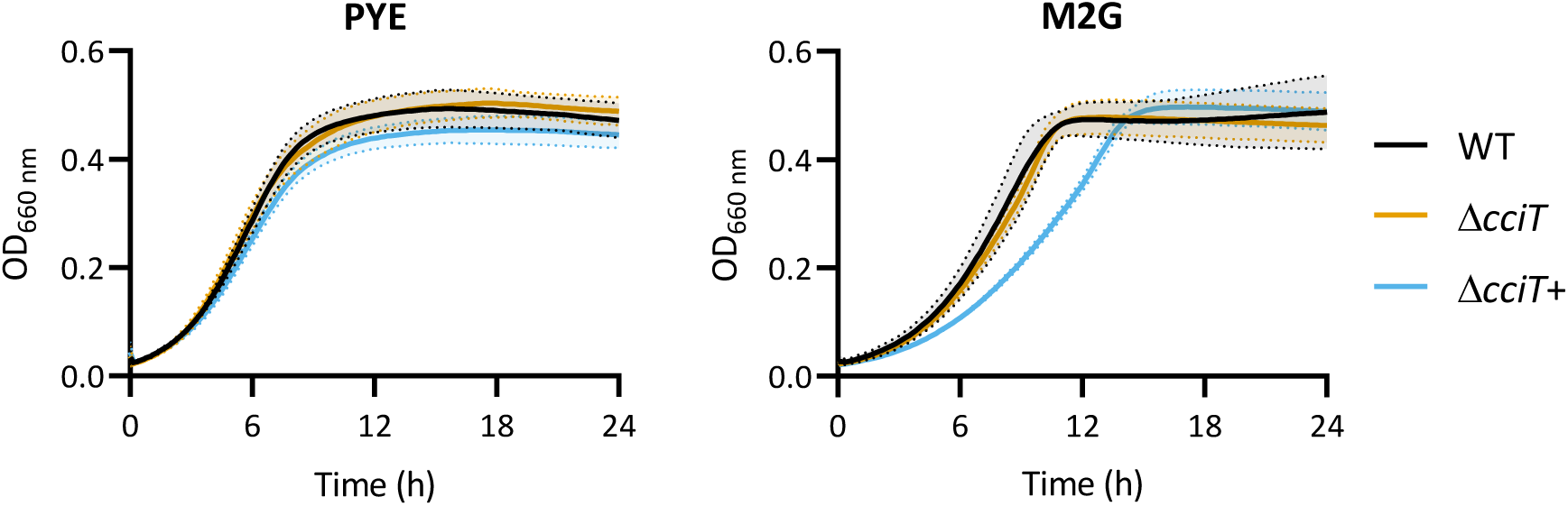
The deletion of *cciT* does not impact bacterial fitness under control conditions. Growth profiles at an absorbance of 660 nm of WT, Δ*cciT*, and Δ*cciT*+ strains in PYE (top) and in M2G (bottom) media. Mean ± SD, at least three biological replicates.

**Figure S2.**
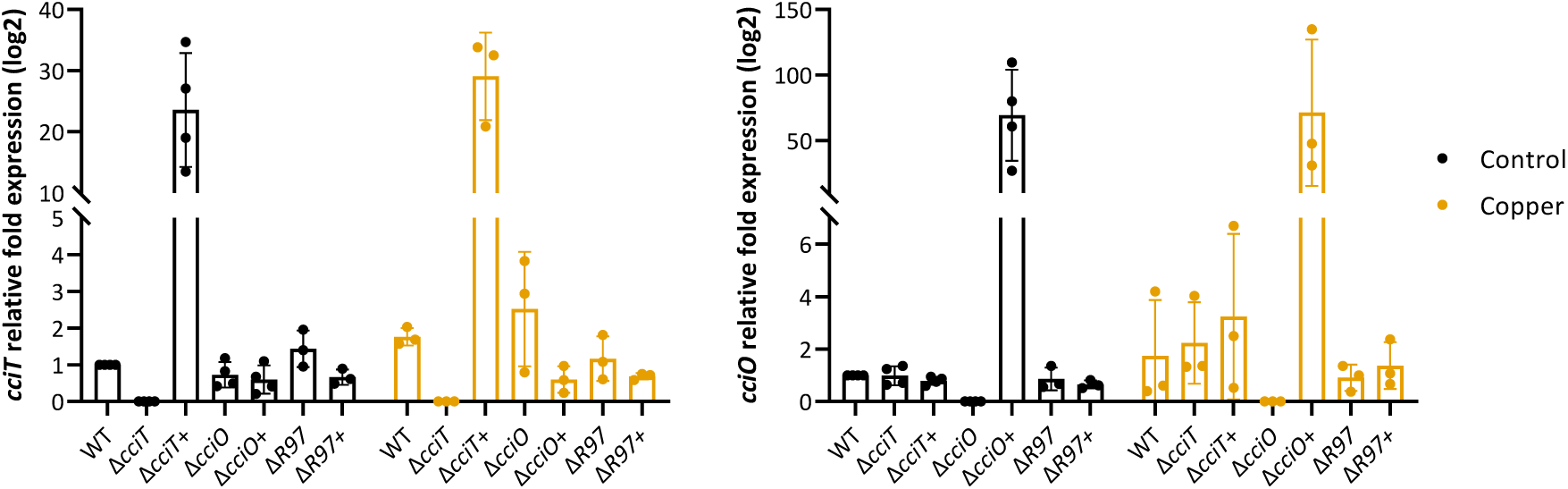
Expression levels of *cciT* and *cciO* genes in the studied genetic backgrounds. Gene expression of *cciT* and *cciO* was measured using RT-qPCR in the studied strains in M2G and after the exposure to CuSO_4_ excess for 10 min. Results are expressed in relative fold changes compared to the WT strain in control condition and calculated as 2^-ΔΔCt^. *mreB* gene has been used as housekeeping gene. Biological replicates ≥ 3, technical replicates = 3, mean ± SD.

**Figure S3.**
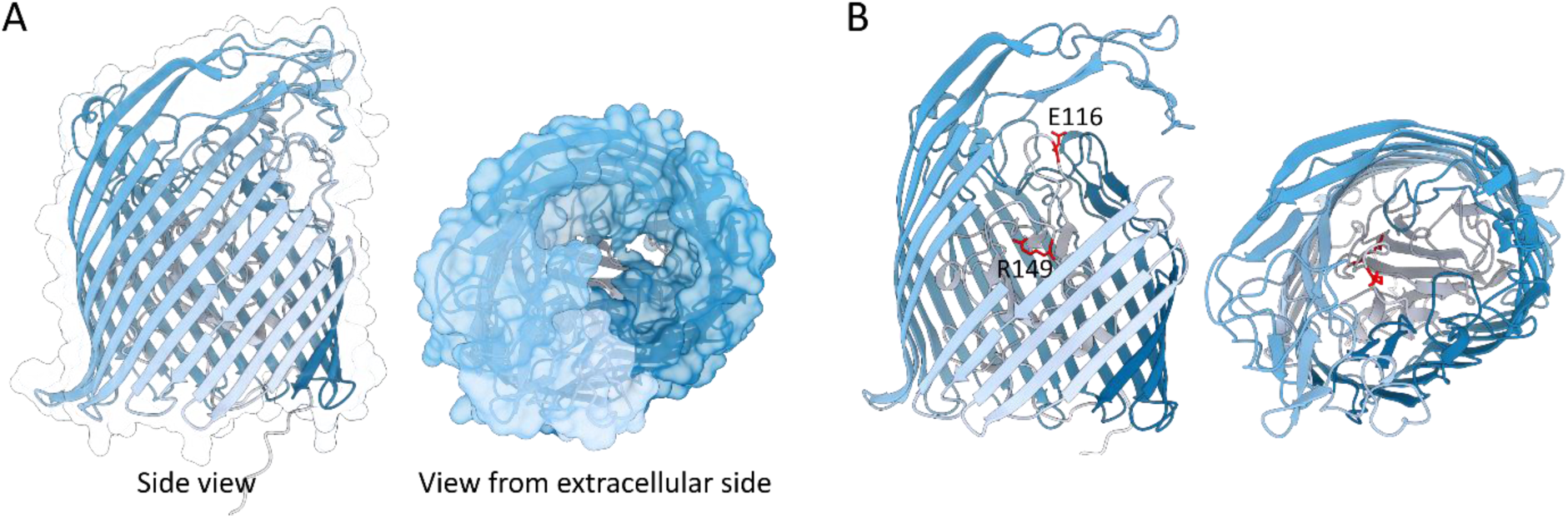
Structure prediction of CciT using AlphaFold. [34]**. A.** TbcT is composed of 22 antiparallel β-strands forming a β-barrel, in which lies the globular plug domain. In white is the potential TonB box. The surface of the β-barrel has been represented, allowing the visualization of the entry of the substrate from the extracellular side. **B.** The residues potentially involved in substrate recognition, E116 and R149, are colored red and the side chains are shown. Models were visualized using ChimeraX [35].

**Figure S4.**
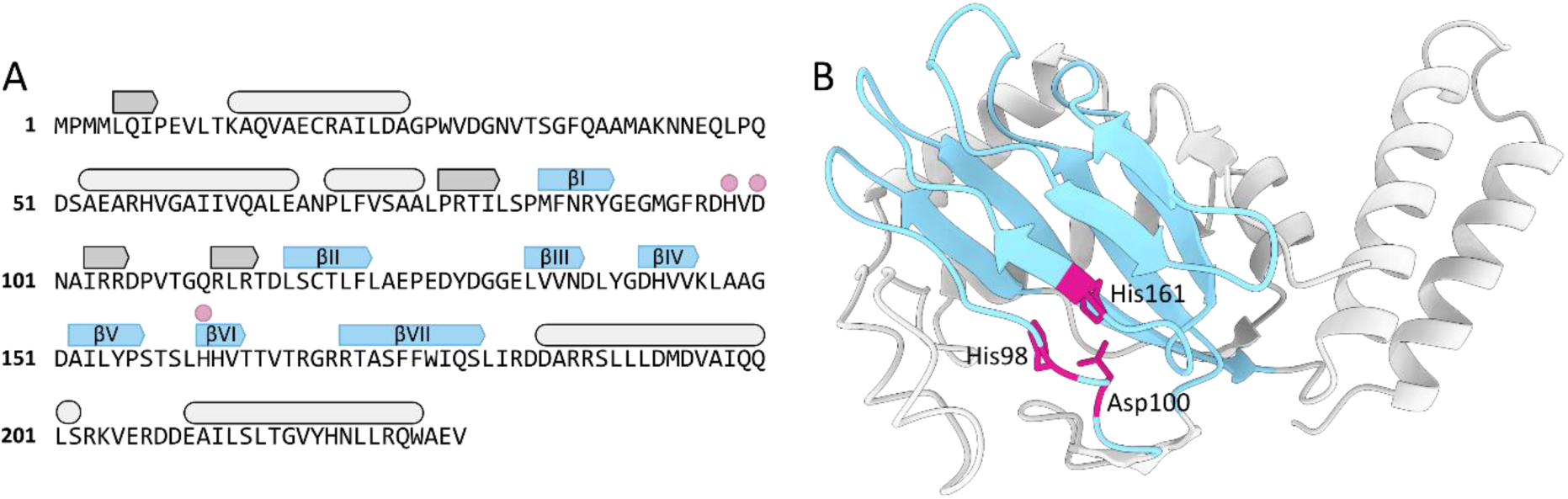
Structure prediction of CciO by AlphaFold. [34]**. A and B.** The DSBH core fold is in light blue and the residues coordinating the Fe^2+^ cofactor are in purple. **A.** Schematic representation of the secondary structure of CciO, based on the 3D structure prediction. β-strands are represented by arrows and α-helices by rounded squares. The additional secondary structures are represented in light grey for α-helices and dark grey for β-strands. **B.** The DSBH fold adopts a squashed barrel conformation, supported by N-terminal secondary structures. The C-terminal α-helices are apart from the core fold. The model was visualized using ChimeraX [35].

